# The abscisic acid could reverse the cuproptosis in sepsis-associated acute kidney injury through the inhibition of ferredoxin-1

**DOI:** 10.1101/2025.06.10.658972

**Authors:** Tianyi Ai, An Shi, Jianxiao Chen, Meishu Ya, Hui Li, Jiajia Tang, Zhiqiang Shi, Zishu Song, Ying Jiang, Mingli Zhu

## Abstract

Sepsis-associated acute kidney injury (S-AKI) is a grave clinical condition with a poor prognosis. Prior studies have observed a potential correlation between copper (Cu) and S-AKI, though the underlying mechanisms remain to be fully elucidated. The present study aims to investigate the underlying mechanism between Cu and renal dysfunction under sepsis conditions. Utilizing a cecal ligation and puncture (CLP) murine model, we observed a substantial upregulation of inflammatory factors and deteriorated renal function (p<0.05), concomitant with an augmented mortality rate (p<0..05). Concurrently, the renal tissue exhibited a series of pathological manifestations, including fibrosis and cell death. Concurrently, the expression of the cuproptosis-regulating protein FDX-1 (p<0.05) was found to be elevated in CLP mice. Collectively, these data indicated the activation of cuproptosis in CLP-induced S-AKI. Intriguingly, the application of abscisic acid (ABA), a FDX-1 inhibitor, led to a substantial alleviation of mortality rate and inflammatory factor level (p<0.05). Furthermore, a significant alleviation was observed in renal pathology and biofunction (p<0.05). Collectively, these observations suggest that ABA exerts a protective effect on renal function during cuproptosis promoted S-AKI. Consequently, it can be concluded that ABA has the potential to reverse cuproptosis in S-AKI by inhibiting FDX-1.

## 1. Introduction

Sepsis is a life-threatening medical emergency caused by an overwhelming inflammatory response to infection. This extreme immune response can lead to significant tissue and organ damage, with the kidneys being the most commonly affected organ^1, 2^. The sepsis-associated acute renal injury (S-AKI) is defined as acute renal failure within seven days after a sepsis attack. Literature demonstrates that the incidence of S-AKI in the intensive care unit (ICU) can reach up to 67%, while the mortality rate ranges from 36.67% to 71.7%^1, 3^. Despite the technological advancements in research and clinical practice, there is currently no alternative treatment or prevention strategy for S-AKI beyond supportive care.

Transition metal ions play a pivotal role in various biochemical processes within living organisms^4^, underscoring their vital function in sustaining life. Metal ion homeostasis is a delicate balance and imbalances in metal ions can lead to pathological changes^4^. Copper (Cu) has been demonstrated to be an essential metal element that affects many biological pathways in human bodies, including energy regulation^5^, mitochondrial respiration^6^, antioxidant^7^. Dysregulated Cu metabolism has been associated with various diseases, including cancer, Wilson’s disease, Menkes disease, and neurodegenerative disorders. Recently, a distinctive form of programmed cell death, designated as cuproptosis, has been identified^6^. This novel form of cell death is distinct from oxidative stress-related cell death (e.g., apoptosis, ferroptosis, and necroptosis) because the process of cuproptosis is initiated when the intracellular concentration of Cu exceeds the homeostatic threshold. Excessive Cu induces mitochondrial stress, which results in the accumulation of fatty acylated mitochondrial enzymes and the loss of Fe–S cluster proteins^6^. The eventual outcome of this process is the induction of cell death, attributable to the deleterious effects of protein toxicity stress within the cytosol. To date, cuproptosis research has been predominantly focused on cancers^8-10^ and cardiovascular disease^11-13^. However, there is a paucity of research investigating the potential toxicological consequences of cuproptosis on kidney injury.

Recent findings have indicated that sepsis plays a pivotal role in the process of cuproptosis^14-16^. This interaction between sepsis and cuproptosis has been observed in various pathological processes, including lung^17, 18^ and cardiomyopathy injury^14, 15^. Although previous studies had demonstrated the toxic effect of Cu on kidney^19, 20^, the role of cuproptosis in kidney injury remains largely unknown. In this regard, we hypothesize that cuproptosis plays a pivotal role in S-AKI and seek to elucidate the underlying mechanisms. Furthermore, previous studies demonstrated that the ferredoxin 1 (FDX-1) could degrade Fe–S cluster proteins and facilitate cuproptosis^6^ while knockout of the *FDX-1* gene attenuated cuproptosis^21^. In light of the well-established role of abscisic acid (ABA) in reducing the expression of FDX-1, this study sought to investigate the potential of ABA to reverse S-AKI in the context of cuproptosis inhibition.

## 2. Method

The present study was approved by the Institutional Review Board (IRB) of the Shanghai Jiaotong University School of Medicine Animal Care and Use Committee and was performed in accordance with the National Institutes of Health guidelines for the care and use of animals in research.

### 2.1. Animal Husbandry

The male C57/BL6 mice were obtained and raised in the Shanghai Jiaotong University Animal Laboratory. The mice were maintained on a 12-hour light/dark cycle, and all mice were fed ad libitum. At six weeks of age, 60 mice were randomly assigned to one of three groups: a control group, a cecal ligation and puncture (CLP) group to mimic a sepsis condition, and a CLP group treated with abscisic acid (CLP/ABA group). The CLP model was created as previously described^22^ while the control group received sham surgery. For the CLP/ABA groups, mice were administered 100 mg/kg of ABA intravenously. Twenty-four hours after sham or CLP surgery, mice were euthanized by cervical dislocation, and blood and renal tissues were collected for further analysis.

### 2.2. H&E staining

The H&E staining procedure was carried out in accordance with the established protocol. The renal tissue was subjected to a series of ethanol concentrations, followed by clearing in xylene and finally embedded in paraffin. Subsequently, the lung was sectioned at a thickness of approximately 4 μm, and these slices were subjected to hematoxylin and eosin staining. Finally, the stained slices were examined under an optical microscope at magnifications ranging from x100 to x400.

### 2.3. Masson’s trichrome staining

The tissue was initially dehydrated through the application of increasingly concentrated ethanol solutions. Subsequently, the tissue was cleared with xylene and infiltrated with paraffin in a paraffin embedding bath. Thereafter, the tissue was cooled and solidified with liquid paraffin. Then, the cooled wax block was fixed, and the thickness of the slices was adjusted to 4 μm to ensure uniformity. The slices were then deparaffinized using xylene, followed by absolute ethanol and alcohol in a step-by-step manner. Weigert’s iron hematoxylin staining was applied to the slices, and the samples were sealed with neutral gum. The samples were then observed and analyzed under an optical microscope.

### 2.4. TUNEL assay

The tissue sections were fixed in paraffin and subsequently treated with proteinase K to facilitate permeabilization. Following the completion of the permeabilization process, the samples were examined using a fluorescence microscope. To label the samples, they were exposed to dUTP, a molecule that is specifically incorporated into DNA by the enzyme TdT. Following the catalysis of dUTP incorporation, the samples were examined under a fluorescence microscope. Under standard fluorescence filter settings, green fluorescence was observed at a wavelength of approximately 520 ± 20 nm. This finding indicated the presence of FITC-12-dUTP, which had been incorporated into the DNA of apoptotic cell nuclei. Concurrently, red fluorescence resulting from propidium iodide (PI) was observed at wavelengths exceeding 620 nm. Blue fluorescence, attributable to 4’,6-diamidino-2-phenylindole (DAPI), was observed at a wavelength of 460 nm. To ensure the preservation of the stained slides, they were stored in darkness at a temperature of 4°C for a period of time. The presence of PI or DAPI in the cell nuclei led to the observation of red or blue staining, respectively, in both apoptotic and non-apoptotic cell nuclei. However, only apoptotic cell nuclei displayed green fluorescence, indicating the specific incorporation of FITC-12-dUTP into their DNA.

### 2.5. Western Blot

Following a 60-second sonication on ice, the tissue mixture was subjected to a 15-minute centrifugation at 12,000 rpm, resulting in the collection of the sediment phase. The bicinchoninic acid (BCA) protein assay was used to determine the protein concentration of sediment. Protein loading buffer was used to quantify protein at a concentration of 5 mg/mL, and it was stored at -20°C for later use. Then the bovine serum albumin standard was diluted in a gradient. Then, various dilutions of protein standards were added to the protein samples, which were then placed in microplates or test tubes. Finally, the BCA working solution was added. The mixture was thoroughly agitated, sealed, and then placed into an incubator set at 37°C. Following this, the samples were allowed to cool to room temperature and the absorbances of the samples were measured at 562 nm or near this wavelength, with a blank serving as the control. Subsequent to this step, the average blank standard’s 562 nm absorption value should be deducted from each standard’s and test protein sample’s 562 nm absorption value. Subsequent to this step, perform SDS-PAGE electrophoresis and electrotransfer the protein bands separated on the gel to a PVDF membrane. The transferred membrane should then be incubated with primary antibodies (1:1000), washed, and mixed with the ECL Kit at a volume ratio of 1:1 (v/v). The mixture should be applied evenly to the surface of the PVDF membrane and incubated for 4 minutes at room temperature. Following this step, the liquid should be removed by shaking, and the membrane should be placed in a chemiluminescence imaging system for imaging.

### 2.6. Statistical analysis

The statistical analyses were conducted using GraphPad Prism 7.0 software and are presented as mean ± standard error of the mean (SEM). The Analysis of Variance (ANOVA) was employed to compare the differences among groups. P-values less than 0.05 were considered statistically significant.

## 3. Result

### 3.1. The CLP mice exhibited signs of acute kidney injury under septic conditions

Twenty-four hours after CLP surgery, a higher mortality rate was observed in the CLP group compared to the control group (p<0.05, Fig 1A). Concurrently, a substantial increase was observed in inflammatory factors [IL-6, TNF-α, IL-1β, and IL-18 (all p<0.01, Fig 1 B-E)]. Collectively, these findings indicated that CLP could induce sepsis in murine models. Furthermore, a significant increase was detected in both serum creatinine (CREA) and blood urea nitrogen (BUN) levels in the CLP mice (both p<0.01, Fig. 1F & G).

**Fig. 1.**
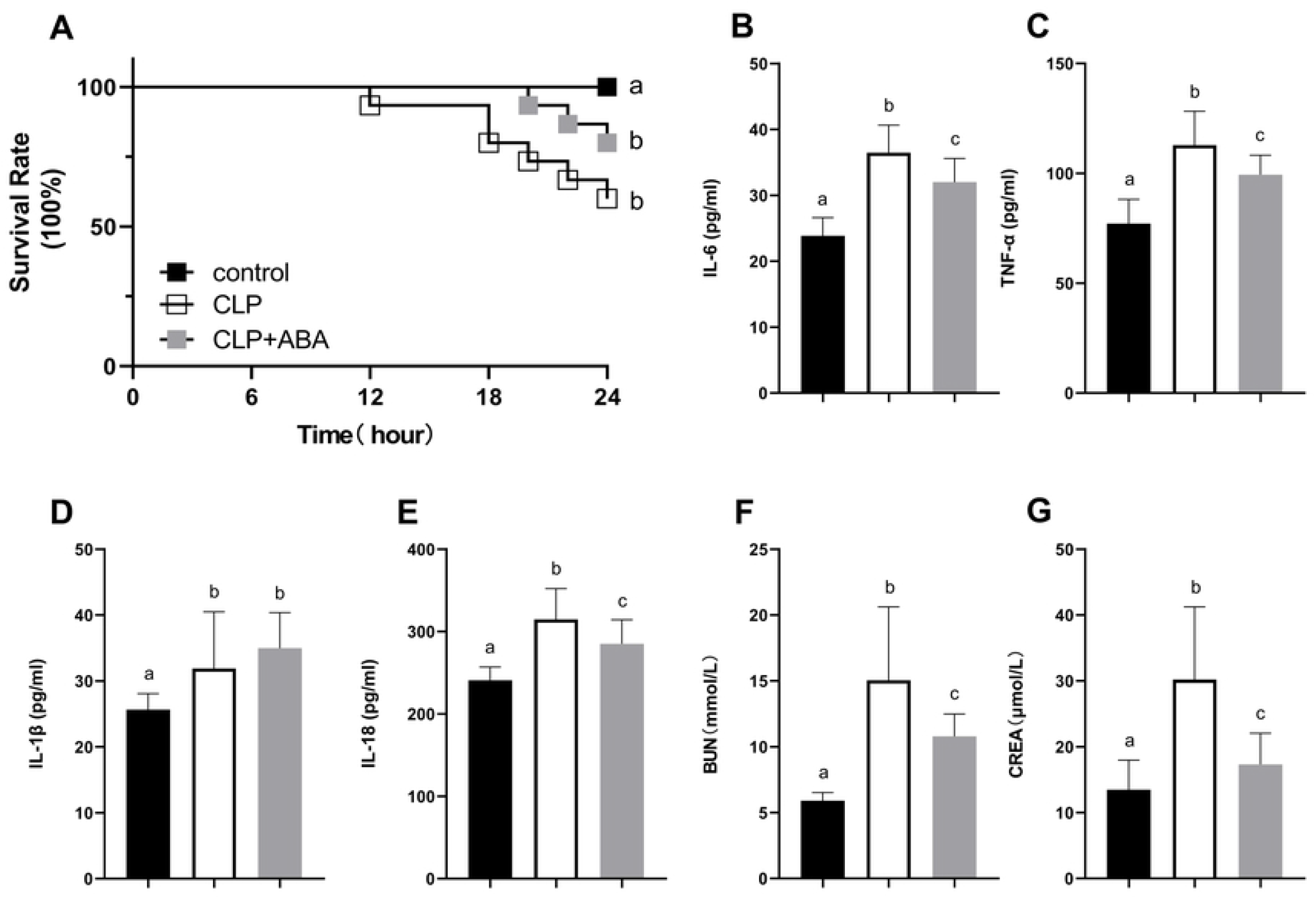
The cecal ligation and puncture (CLP) group of mice exhibited a heightened response to sepsis and a heightened susceptibility to sepsis-associated acute kidney injury (S-AKI). This response could be mitigated by the administration of abscisic acid (ABA). (A) Twenty-four hours after CLP surgery, the CLP group exhibited a significantly elevated mortality rate compared to the control group (p < 0.01). Concurrently, a substantial increase was observed in four tested inflammatory factors, including IL-6, TNF-α, IL-1β, and IL-18 (all p < 0.01) (B-E) as well as the serum creatinine (CREA) and blood urea nitrogen (BUN) levels (F-G) (both p < 0.05). Subsequent to the application of ABA, a significant decline in mouse mortality was observed (A), accompanied by a decrease in inflammatory factors (B-E) and an improvement in renal function (F-G). *A significant difference is indicated by the presence of different letters in the histogram column.

Consistent with the sepsis and renal function data, mice from the CLP groups exhibited a range of pathological changes in the kidney, including vacuolar degeneration of renal tubular epithelial cells, interstitial edema, sloughing of renal tubular epithelial cells, dilation of the renal tubular lumen (Fig. 2C & D), and fibrosis (Fig. 3), accompanied by an escalation of cell death (Fig. 4). Collectively, these findings demonstrate that CLP could significantly exacerbate sepsis, potentially leading to S-AKI in mice.

**Fig. 2.**
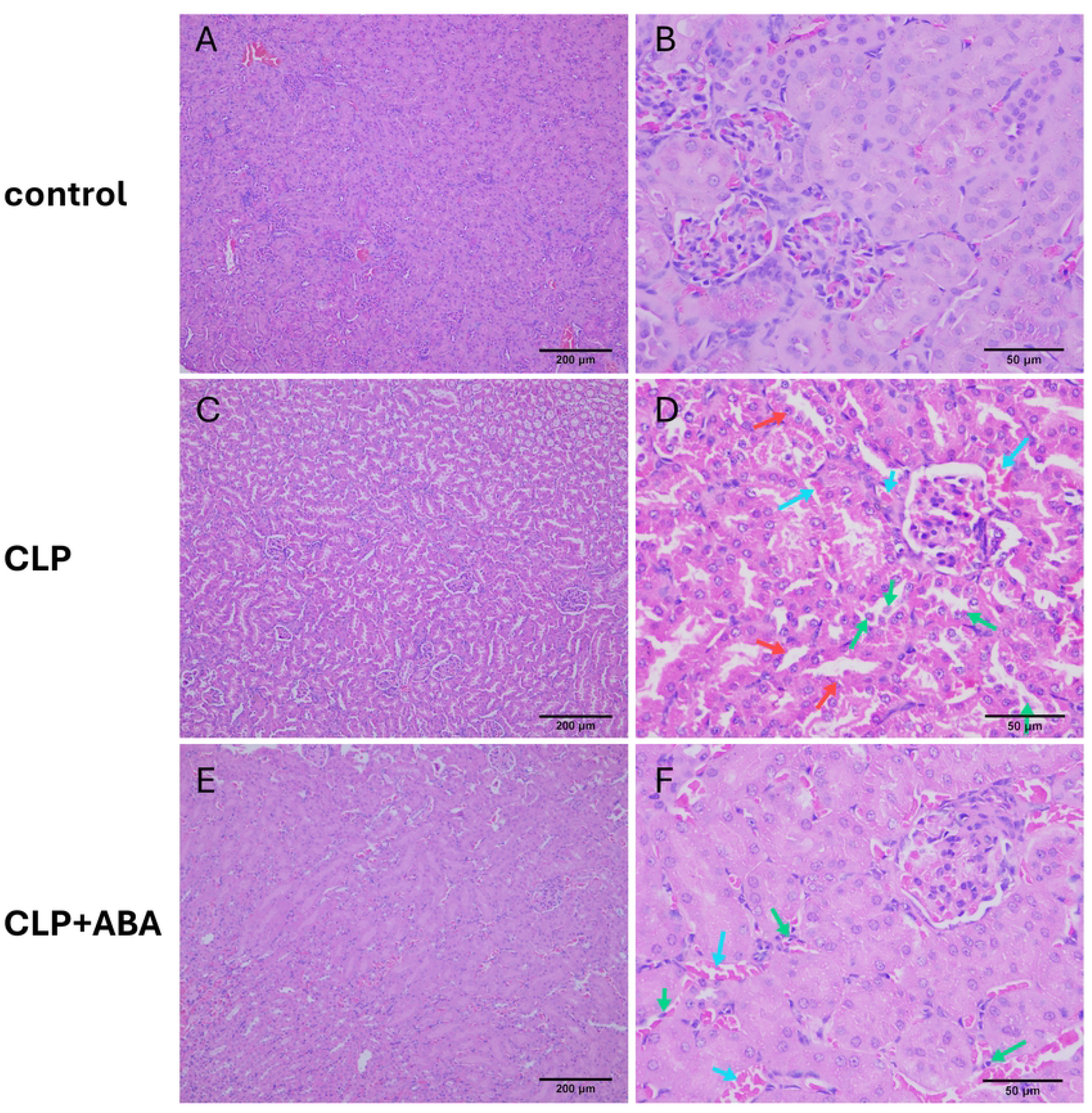
The cecal ligation and puncture (CLP) model has been demonstrated to induce sepsis-associated acute kidney injury (S-AKI) with a kidney pathology profile that can be mitigated by abscisic acid (ABA). In the control group, renal H&E samples exhibited normal renal structure (A & B). However, following CLP, a range of pathological changes were observed in the kidneys of the mice on H&E staining, including vacuolar degeneration of renal tubular epithelial cells, interstitial edema, sloughing of renal tubular epithelial cells, and dilation of the renal tubular lumen (C & D). Following the administration of ABA, these pathological alterations were mitigated (E & F).

**Fig. 3.**
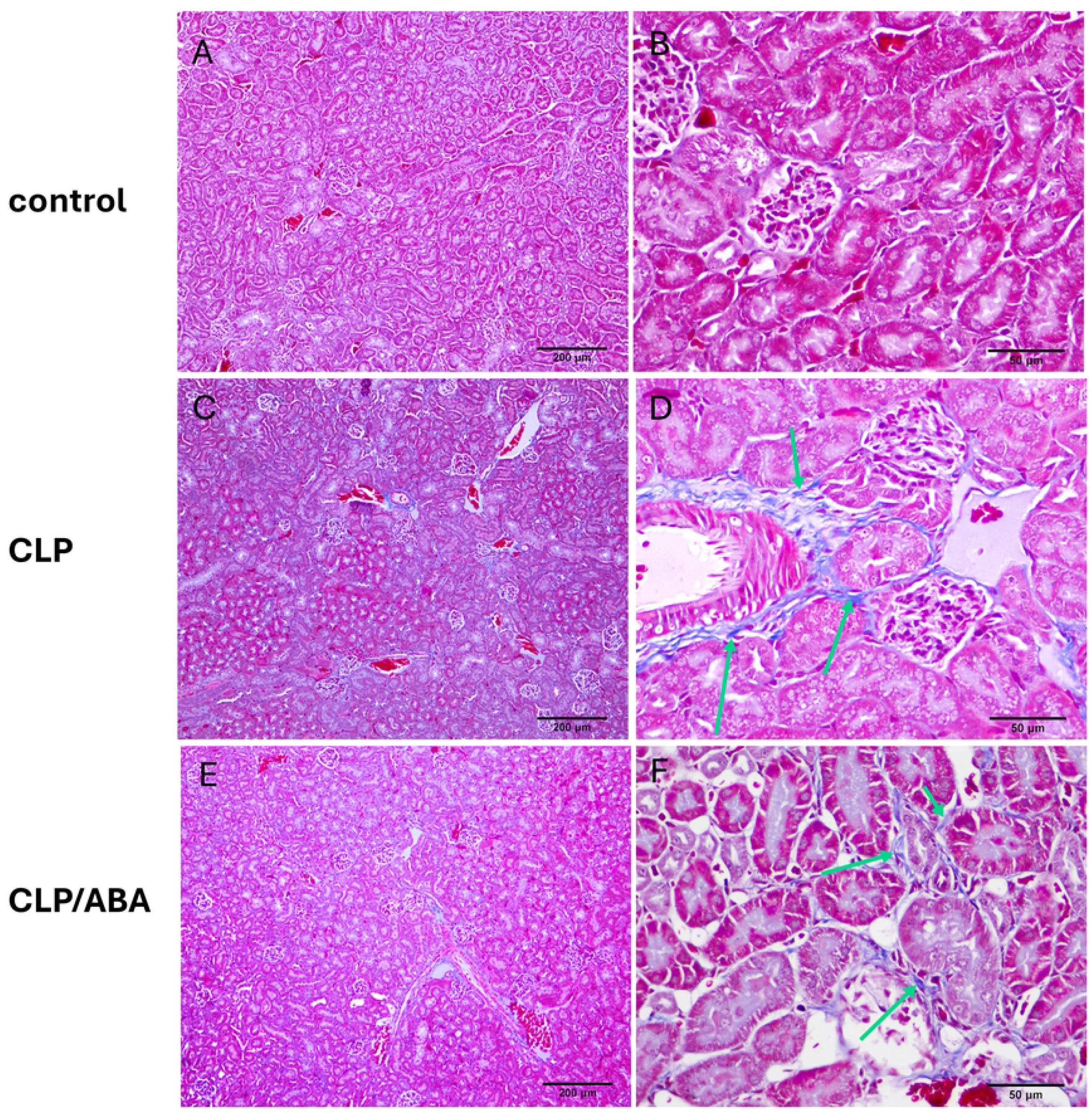
The cecal ligation and puncture (CLP) model has been demonstrated to induce sepsis-associated acute kidney injury (S-AKI), characterized by kidney fibrosis. This phenomenon can be mitigated by the application of abscisic acid (ABA). The Masson’s trichrome staining revealed a significant increase in fibrosis in the renal tissue (C & D) compared to the control group (A & B). Following the administration of ABA, a significant mitigation of fibrosis was observed (E & F).

**Fig. 4.**
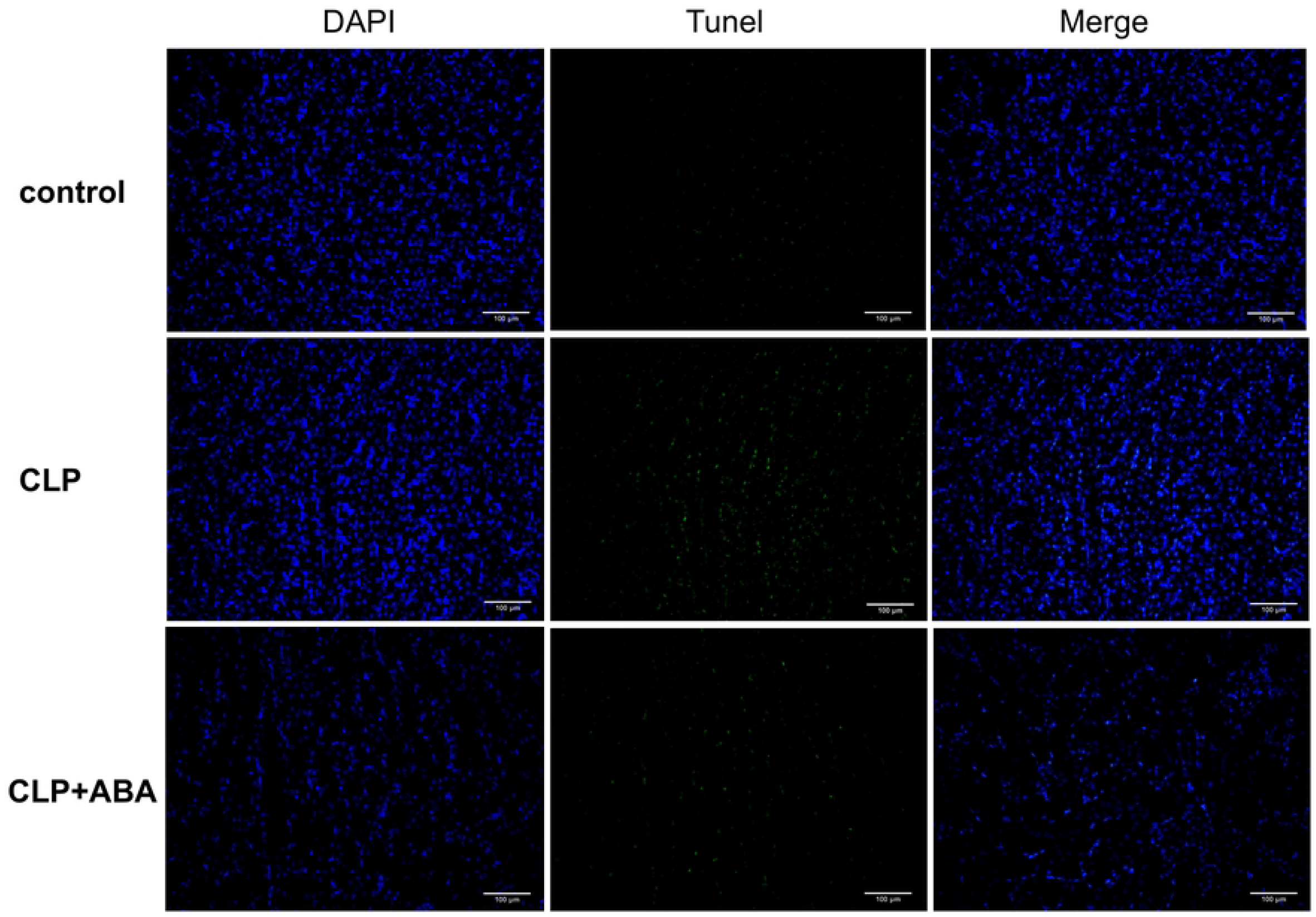
The cecal ligation and puncture (CLP) model has been demonstrated to induce kidney cell death, which can be mitigated by the application of abscisic acid (ABA). The TUNEL assay revealed a significant increase in cell death in the renal tissue compared to the control group. However, the administration of ABA led to a substantial mitigation of cell death.

### 3.2. The cuproptosis is implicated in the occurrence of CLP-induced S-AKI

In accordance with the prevailing hypotheses, the CLP mice exhibited significant upregulation of FDX-1 in renal tissue (p<0.01) (Fig. 5). Given the prior identification of the FDX-1 as a positive contributor to the cuproptosis^6^, this phenomenon suggests a robust correlation between CLP-induced S-AKI and cuproptosis.

**Fig. 5.**
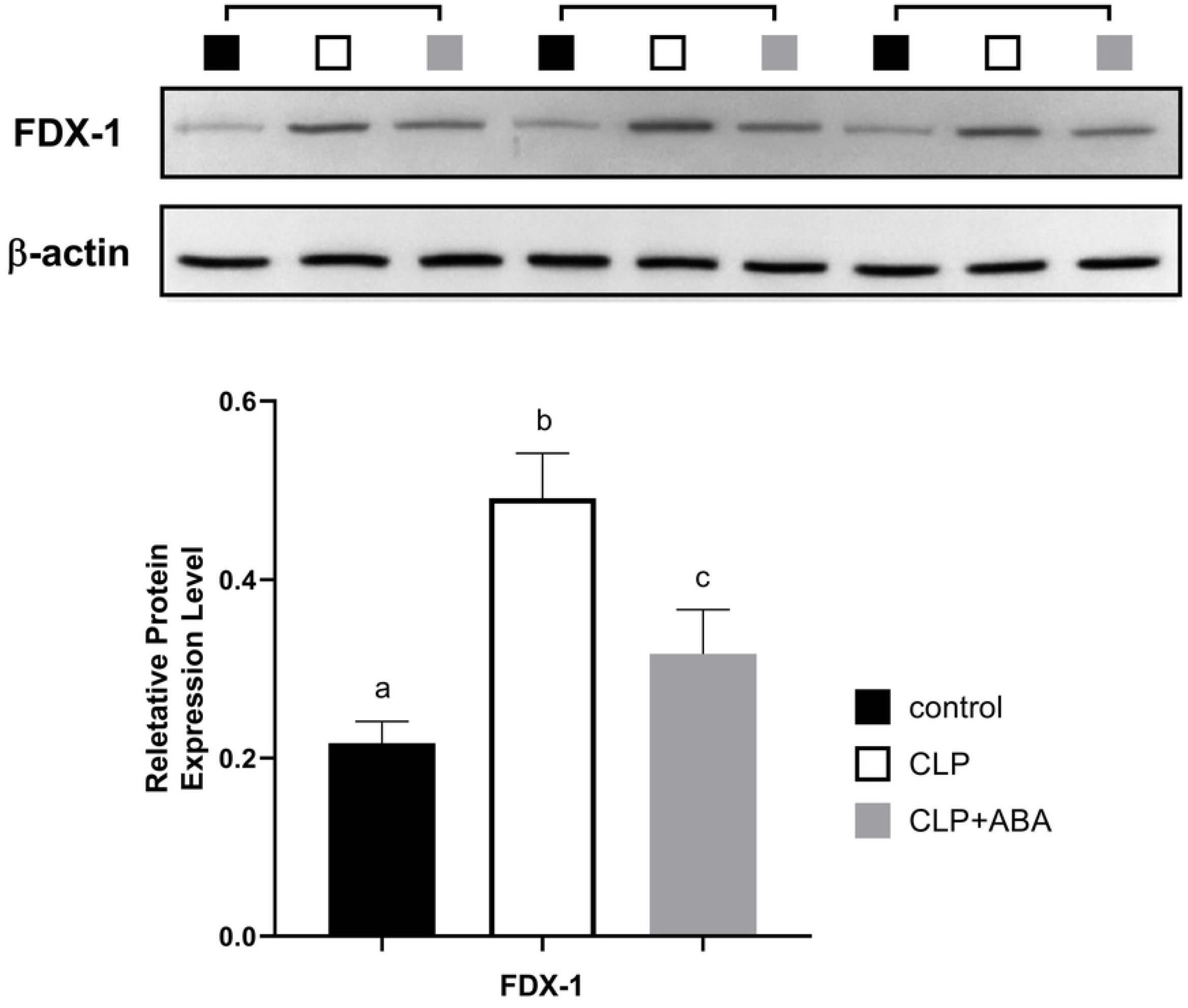
The cecal ligation and puncture (CLP) mice exhibited significant upregulation of FDX-1 in renal tissue, suggesting the activation of cuproptosis. However, the application of abscisic acid (ABA) led to a substantial inhibition of FDX-1 expression, indicating a potential alleviation of cuproptosis.

### 3.3. ABA could alleviate the S-AKI syndrome induced by CLP

The ABA was previously shown to impede FDX-1 expression via binding to *FDX-1* promoter and suppress its transcription^23^. Thus, we hypothesized that the administration of ABA could mitigate the CLP-induced S-AKI by suppressing FDX-1. Our observations revealed that the ABA could lower the mortality in CLP mice (Fig. 1A). This increased survival could be related to alleviated inflammation in the CLP/ABA group, as a significant drop of IL-6, TNF-α, and IL-18 were observed (all p<0.05, Fig. 1B-E). Meanwhile, dropped BUN and CREA also demonstrated relieved renal function by ABA after CLP (both p<0.05, Fig. 1F & G). In comparison with the CLP group, the CLP+ABA mice exhibited only mild vacuolar degeneration of renal tubular epithelial cells and interstitial edema (Fig. 2E & F), accompanied by a reduction in renal fibrosis (Fig. 3E & F) and cell death (Fig. 4). Moreover, the ABA significantly inhibit the FDX-1 expression after CLP treatment (p<0.05, Fig. 5). Consequently, these observations collectively indicated that the role of ABA in attenuating cuproptosis process, which restored partial renal function and thus increased survival rate.

## 4. Discussion

Cu is an essential element in virtually all living organisms. Numerous studies have demonstrated that Cu serves as a cofactor for a variety of key metabolic enzymes that drive a broad range of physiological processes. Thus, failure to maintain the Cu concentration could disrupt various biochemical processes and subsequent novel regulatory mechanisms of cell death called cuproptosis. Previous studies have demonstrated the significant role of cuproptosis in the pathogenesis of various kidney diseases, chronic kidney disease, diabetic nephropathy, kidney transplantation, and kidney stones. However, studies on the role of cuproptosis in S-AKI are rare. In the present study, we demonstrated that cuproptosis plays a pivotal role in CLP-induced S-AKI, as indicated by its hallmarks of renal pathological and functional changes, as well as elevated FDX-1 expression. Conversely, administration of ABA has been observed to mitigate cuproptosis and offer protection against S-AKI in murine models. This study is the first to demonstrate the critical role of cuproptosis in S-AKI and the potential for ABA administration to alleviate this condition.

Mitochondria is the central site of oxidative metabolism. As a mitochondrial protein, the FDX has long been identified as a family of redox proteins involving in various redox process^24^. For FDX-1, it transfers electrons from NADPH and donates electrons to cytochrome p450 systems inside the mitochondria, which enlightens its crucial role in multiple metabolic processes^24^. Interestingly, recent studies also revealed the important role of FDX-1 in cuproptosis via two mechanisms. First, FDX-1 could act as a reductase to reduce Cu^+2^ to its more toxic form Cu^+1 25^. Second, FDX-1 could lipoylate several tricarboxylic acid (TCA) cycle enzymes and cause abnormal acylated protein aggregation inside mitochonria^6^. Taken together, FDX-1 becomes the key factor during cuproptosis. In the current study, we observed that sepsis could lead to elevation of FDX-1 expression in the renal tissue, which was concurrent with clinical manifestation of S-AKI in septic mice both histologically and biochemically. Although we did not measure the lipid metabolism and [Cu^+1^] changes inside the renal tissue, the present findings indicate that the induction of cuproptosis in renal tissue may serve as a potential underlying mechanism contributing to the development of S-AKI.

The ABA is a phytohormone that plays a vital role in the growth and regulation of different processes in plants. Particularly, the ABA plays an important role in plant adaptation to both biotic and abiotic stress^26, 27^, which is mainly due to its regulatory function in abundant gene transcription and RNA processing^28^. A recent study demonstrated that ABA can reduce the expression of FDX-1 through its binding to the FDX-1 promoter in conjunction with the transcription factor ABI5^23^. Given the significant elevation of FDX-1 observed in the S-AKI model, we investigated the potential function of ABA in reversing or mitigating the process of cuproptosis. Subsequent to the administration of ABA to mice, the renal tissue exhibited alleviated pathology and biochemical biomarkers, which were concomitant with an augmented survival rate of S-AKI mice. Consequently, ABA could impede the cuproptosis process via FDX-1 inhibition.

The present study is subject to several limitations. First, it investigates FDX-1-mediated cuproptosis in sepsis-related acute renal injury but lacks further explanation and exploration of the molecular mechanisms involved (for example, the [Cu^+1^], [Cu^+2^], Fe-S cluster protein, etc). Additional experiments, both in vivo and in vitro, are necessary to determine whether knocking out the FDX-1 gene can reverse the organ damage caused by sepsis AKI. Secondly, while the findings demonstrate the potential of ABA to impede copper-induced cell death and mitigate renal injury, further research is required to elucidate its underlying mechanisms in sepsis.

## 5. Conclusion

In summary, our research has determined that sepsis can induce S-AKI through the process of cuproptosis activation and FDX-1 is a critical biomarker during this process. Following administration of ABA, we observed a decrease in FDX-1 expression, which counteracted the pathological outcomes associated with cuproptosis in the renal tissue. Given the critical nature of S-AKI, the present study offers significant insights into the role of cuproptosis in this condition and suggests potential therapeutic interventions for S-AKI during clinical practice.

## Acknowledgements

We would like to thank Jingli Guo for experimental guidance.

## Notes

### Competing Interest Statement

The authors have declared no competing interest.

## Reference

1. Liu J, Xie H, Ye Z, et al. Rates, predictors, and mortality of sepsis-associated acute kidney injury: a systematic review and meta-analysis. BMC nephrology 2020; 21: 318. 2020/08/02. DOI: 10.1186/s12882-020-01974-8.

2. Alobaidi R, Basu RK, Goldstein SL, et al. Sepsis-associated acute kidney injury. Seminars in nephrology 2015; 35: 2–11. 2015/03/22. DOI: 10.1016/j.semnephrol.2015.01.002.

3. Hsu YC and Hsu CW. Septic acute kidney injury patients in emergency department: The risk factors and its correlation to serum lactate. The American journal of emergency medicine 2019; 37: 204–208. 2018/05/20. DOI: 10.1016/j.ajem.2018.05.012.

4. Valko M, Jomova K, Rhodes CJ, et al. Redox- and non-redox-metal-induced formation of free radicals and their role in human disease. Archives of toxicology 2016; 90: 1–37. 2015/09/08. DOI: 10.1007/s00204-015-1579-5.

5. Guo H, Wang Y, Cui H, et al. Copper Induces Spleen Damage Through Modulation of Oxidative Stress, Apoptosis, DNA Damage, and Inflammation. Biological trace element research 2022; 200: 669–677. 2021/03/20. DOI: 10.1007/s12011-021-02672-8.

6. Tsvetkov P, Coy S, Petrova B, et al. Copper induces cell death by targeting lipoylated TCA cycle proteins. Science (New York, NY) 2022; 375: 1254–1261. 2022/03/18. DOI: 10.1126/science.abf0529.

7. Jian Z, Guo H, Liu H, et al. Oxidative stress, apoptosis and inflammatory responses involved in copper-induced pulmonary toxicity in mice. Aging 2020; 12: 16867–16886. 2020/09/22. DOI: 10.18632/aging.103585.

8. Wang W, Lu K, Jiang X, et al. Ferroptosis inducers enhanced cuproptosis induced by copper ionophores in primary liver cancer. Journal of experimental & clinical cancer research : CR 2023; 42: 142. 2023/06/06. DOI: 10.1186/s13046-023-02720-2.

9. Sun L, Zhang Y, Yang B, et al. Lactylation of METTL16 promotes cuproptosis via m(6)A-modification on FDX1 mRNA in gastric cancer. Nature communications 2023; 14: 6523. 2023/10/21. DOI: 10.1038/s41467-023-42025-8.

10. Qin Y, Liu Y, Xiang X, et al. Cuproptosis correlates with immunosuppressive tumor microenvironment based on pan-cancer multiomics and single-cell sequencing analysis. Molecular cancer 2023; 22: 59. 2023/03/25. DOI: 10.1186/s12943-023-01752-8.

11. Wang D, Tian Z, Zhang P, et al. The molecular mechanisms of cuproptosis and its relevance to cardiovascular disease. Biomedicine & pharmacotherapy = Biomedecine & pharmacotherapie 2023; 163: 114830. 2023/05/08. DOI: 10.1016/j.biopha.2023.114830.

12. Kunutsor SK, Dey RS and Laukkanen JA. Circulating Serum Copper Is Associated with Atherosclerotic Cardiovascular Disease, but Not Venous Thromboembolism: A Prospective Cohort Study. Pulse (Basel, Switzerland) 2021; 9: 109–115. 2022/01/28. DOI: 10.1159/000519906.

13. Sudhahar V, Das A, Horimatsu T, et al. Copper Transporter ATP7A (Copper-Transporting P-Type ATPase/Menkes ATPase) Limits Vascular Inflammation and Aortic Aneurysm Development: Role of MicroRNA-125b. Arteriosclerosis, thrombosis, and vascular biology 2019; 39: 2320–2337. 2019/09/27. DOI: 10.1161/atvbaha.119.313374.

14. Yan J, Li Z, Li Y, et al. Sepsis induced cardiotoxicity by promoting cardiomyocyte cuproptosis. Biochemical and biophysical research communications 2024; 690: 149245. DOI: 10.1016/j.bbrc.2023.149245.

15. Song J, Ren K, Zhang D, et al. A novel signature combing cuproptosis- and ferroptosis-related genes in sepsis-induced cardiomyopathy. Frontiers in genetics 2023; 14: 1170737. 2023/04/11. DOI: 10.3389/fgene.2023.1170737.

16. Wang Y, Qiu X, Liu J, et al. Cuproptosis-Related Biomarkers and Characterization of Immune Infiltration in Sepsis. Journal of inflammation research 2024; 17: 2459–2478. 2024/04/29. DOI: 10.2147/jir.S452980.

17. Wang Y, Zhao Z and Xiao Z. The Emerging Roles of Ferroptosis in Pathophysiology and Treatment of Acute Lung Injury. Journal of inflammation research 2023; 16: 4073–4085. 2023/09/20. DOI: 10.2147/jir.S420676.

18. Zhu M, Tang X, Xu J, et al. Cuproptosis-related Gene Signatures and Immunological Characterization in Sepsis-associated Acute Lung Injury. Combinatorial chemistry & high throughput screening 2024 2024/05/23. DOI: 10.2174/0113862073290692240509094709.

19. Jiayi H, Ziyuan T, Tianhua X, et al. Copper homeostasis in chronic kidney disease and its crosstalk with ferroptosis. Pharmacological research 2024; 202: 107139. 2024/03/15. DOI: 10.1016/j.phrs.2024.107139.

20. Li Y, Zhong G, He T, et al. Effect of arsenic and copper in kidney of mice: Crosstalk between Nrf2/Keap1 pathway in apoptosis and pyroptosis. Ecotoxicology and environmental safety 2023; 266: 115542. 2023/10/07. DOI: 10.1016/j.ecoenv.2023.115542.

21. Wang J, Luo LZ, Liang DM, et al. Progress in the research of cuproptosis and possible targets for cancer therapy. World journal of clinical oncology 2023; 14: 324–334. 2023/09/29. DOI: 10.5306/wjco.v14.i9.324.

22. Dejager L, Pinheiro I, Dejonckheere E, et al. Cecal ligation and puncture: the gold standard model for polymicrobial sepsis? Trends in microbiology 2011; 19: 198–208. 2011/02/08. DOI: 10.1016/j.tim.2011.01.001.

23. Cui W, Wang S, Han K, et al. Ferredoxin 1 is downregulated by the accumulation of abscisic acid in an ABI5-dependent manner to facilitate rice stripe virus infection in Nicotiana benthamiana and rice. The Plant journal : for cell and molecular biology 2021; 107: 1183–1197. 2021/06/22. DOI: 10.1111/tpj.15377.

24. Schulz V, Basu S, Freibert SA, et al. Functional spectrum and specificity of mitochondrial ferredoxins FDX1 and FDX2. Nature chemical biology 2023; 19: 206–217. 2022/10/26. DOI: 10.1038/s41589-022-01159-4.

25. Xue Q, Kang R, Klionsky DJ, et al. Copper metabolism in cell death and autophagy. Autophagy 2023; 19: 2175–2195. 2023/04/15. DOI: 10.1080/15548627.2023.2200554.

26. Zhu JK. Salt and drought stress signal transduction in plants. Annual review of plant biology 2002; 53: 247–273. 2002/09/12. DOI: 10.1146/annurev.arplant.53.091401.143329.

27. Yoshida T, Mogami J and Yamaguchi-Shinozaki K. ABA-dependent and ABA-independent signaling in response to osmotic stress in plants. Current opinion in plant biology 2014; 21: 133–139. 2014/08/12. DOI: 10.1016/j.pbi.2014.07.009.

28. Cutler SR, Rodriguez PL, Finkelstein RR, et al. Abscisic acid: emergence of a core signaling network. Annual review of plant biology 2010; 61: 651–679. 2010/03/03. DOI: 10.1146/annurev-arplant-042809-112122.

